# In Vivo 4D Oxy-Wavelet MRI as a Non-Invasive Biomarker of Brain Mitochondrial Function across the Lifespan

**DOI:** 10.64898/2026.05.21.726892

**Authors:** Devin Raine Everaldo Cortes, Sean Hartwick, Thomas Becker-Szurszewski, Kristina Elsa Schwab, Cody Ruck, Shanim Manzoor, Noah William Coulson, Dalton West, Margaret Caroline Stapleton, Samuel Wyman, Cecilia Wen-Ya Lo, Sivakama Bharathi, Eric S. Goetzman, Anthony G. Christodoulou, Yijen Lin Wu

**Affiliations:** Department of Pediatrics, School of Medicine, University of Pittsburgh, Pittsburgh, PA, USA; Department of Bioengineering, Swanson School of Engineering, University of Pittsburgh, Pittsburgh, PA, USA; Animal Imaging Core, Rangos Research Center at UPMC Children’s Hospital of Pittsburgh, Pittsburgh, PA, USA; Department of Radiological Sciences, David Geffen School of Medicine, at UCLA, Los Angeles, CA, USA; Department of Bioengineering, University of California Los Angeles, Los Angeles, CA, USA

**Keywords:** Mitochondria, brain, fetal, fMRI, low-rank reconstruction, wavelet analysis

## Abstract

Mitochondria are essential for cellular energy production and are particularly critical for brain development and function. Neurons rely predominantly on oxidative phosphorylation for energy production, rendering the brain highly vulnerable to mitochondrial dysfunction. Consequently, impaired mitochondrial function contributes to a broad spectrum of neurological and systemic disorders, making mitochondria attractive therapeutic targets. Despite this importance, there is currently no non-invasive, spatially resolved method to assess mitochondrial function in the intact living brain.

Here, we establish a non-invasive functional MRI approach—4D Oxy-wavelet MRI—to probe in vivo mitochondrial electron transport chain (ETC) function in a spatially specific manner across the lifespan, from fetal to adult brains. This method employs a low-rank *k*-t sub-Nyquist acquisition strategy to achieve simultaneous structural and functional imaging with high spatial (78 μm) and temporal (∼14 ms) resolution, enabling motion-robust imaging in multi-fetal mouse pregnancies. Mitochondrial ETC function is interrogated by measuring oxygen homeostasis responses to brief hypoxic challenges, analyzed using computational time-frequency wavelet profiling.

We validate this approach in mouse models of mitochondrial respiratory chain disease and late-onset Alzheimer’s disease, from *in utero* fetuses to adults, and demonstrate reproducibility and specificity using pharmacological hyperemia and ETC complex I inhibition. We further show parallel wavelet responses in placenta and fetal brain, enabling multi-organ interrogation of the placenta–brain axis. Finally, we present first-in-human feasibility data, supporting translational potential for non-invasive assessment of mitochondrial function in living brains across the lifespan.

## Introduction

Mitochondria generate most of the cellular energy required for survival and function, especially in highly metabolically active tissues such as the brain. Proper mitochondrial function is critical for neuronal health, and even subtle disruptions can lead to severe, widespread neurological consequences. Mitochondrial dysfunction is a critical contributor to diverse inborn and acquired brain pathologies^1–5^. Neurons are highly sensitive to changes in mitochondrial metabolism because oxidative phosphorylation (OxPhos), not glycolysis, powers synaptic activity, and broadly neurodevelopment, neurogenesis^7–9^, and synaptic plasticity^10^. Furthermore, mitochondrial function is associated with epigenetic regulations^11–15^, influencing biological processes, human diseases, and neurodevelopment. Mitochondria are therefore emerging therapeutic targets^16–27^.

Despite their central role in brain health, no non-invasive method is available to probe *in vivo* mitochondrial functions in the intact brain in a spatially resolved manner. Regional delineation is critical because cognitive impairments resulting from brain disorders manifest in a region-specific manner, yet current approaches do not provide spatially resolved functional mitochondrial readouts for diagnosis or therapeutic monitoring. This study establishes a non-invasive functional magnetic resonance imaging (fMRI) methodology that can probe *in vivo* brain mitochondrial ETC functions with high spatial resolution across the lifespans, from fetuses to adults.

Mouse models are indispensable given the ethical and technical challenges of human pregnancy studies. Mice offer key advantages: each pregnancy carries 6–12 fetuses with different genotypes in the same maternal environment, enabling separation of maternal versus fetal genetic effects. However, in utero multi-fetal mouse fMRI is extremely challenging due to unpredictable fetal motion from multiple heartbeats, and small fetal size. Random fetal orientations further prevent the standard strategy of aligning low-resolution fMRI planes to a high-resolution anatomical template.

This manuscript introduces and evaluates a 4D “Oxy-wavelet MRI” method to probe *in vivo* brain mitochondrial functions in multi-fetal mouse pregnancy to adult brains with both high spatial (78 μm) and temporal (∼14 msec) resolutions. This method uses low-rank *k*-t space-sampling schema that allow sub-Nyquist acquisition for simultaneous structural and functional imaging. The rapid resolution capabilities capture physiological changes for probing ETC function and resolves multi-fetal motion. Leveraging the requirement of mitochondrial ETC activity for maintaining cellular oxygen homeostasis during acute hypoxia, the method uses short hypoxia challenges to probe *in vivo* mitochondrial responses. *In vivo* mitochondrial function is evaluated in MRCD and LOAD mouse models, from multi-fetal mouse brains to adults, using computational time–frequency wavelet profiling of BOLD signals. Reproducibility and specificity are assessed with pharmacological modulation of hyperemia and complex I inhibition.

Placental–fetal interactions profoundly influence fetal neurodevelopment, forming the “*placenta–brain axis*”^28–31^. Maternal oxygen and nutrient delivery critically shape fetal growth and neurodevelopment^32,33^. The placenta is also a neuroendocrine organ^34^, providing neurohormones and signaling molecules crucial for fetal brain development^35^. Thus, the placenta is referred to as “the third brain”^36,37^. Thus, we investigated whether the wavelet responses in the placenta parallel the fetal brain and thus whether -4D Oxy-wavelet MRI can be applied to probe the placenta-brain axis. Finally, as 4D Oxy-wavelet MRI is non-invasive, it is promising for translation to humans. Initial human acquisition demonstrates feasibility for probing mitochondrial function in live brains across the lifespan.

## Results

### 4D *in utero* Oxy-wavelet MRI for simultaneous structural and functional multi-fetal imaging without motion artifact

A novel non-invasive 4D Oxy-wavelet MRI was developed to capture anatomical and functional features in a single scan with high-spatial (isotropic 78μm) and temporal resolution (frame rate: ∼14 msec) using a sub-Nyquist sparse sampling scheme. It allows simultaneous *in utero* brain and placental <u>anatomical</u> and <u>functional</u> imaging of multiple mouse fetuses in a single 4D scan without motion artifact (Fig.1). Mouse fetuses are very small (Fig.1A). Crown-rump length (CRL) for mouse fetuses^38–40^ on the embryonic day E16.5 (Fig. 1A top) and E14.5 (Fig.1A bottom) is ∼16.2 mm and 9.1 mm respectively. Conventional fast imaging, such as echo planar imaging (EPI), which rapidly reverses the readout or frequency encoding-gradient for fast *k*-space sampling still produces motion artifacts in multi-fetal pregnancy (Fig.1B). Our method allows free-breathing-gating-free 4D imaging without motion artifacts (Fig.1C-H), overcoming maternal respiration and multi-fetal motion. Our low-rank *k*-t space sampling scheme expresses dynamic blood oxygenation level dependent (BOLD) signal *ρ*(𝐫, *t*) (for spatial position 𝐫 and time *t*) as the product of basis images 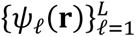 (Fig.1C orange) and temporal functions *φ*_ℓ_(*t*) (Fig.1C blue): 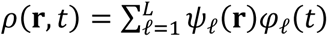. It further exploits the transform sparsity of the image series to allow 4D acquisition (Fig.1C) and reconstruction (Fig.1D) with high spatiotemporal resolutions in one scan^41–43^. The 4D MRI yields a time series of 3D isotropic stacks (Fig.1E), allowing reorientation to visualize individual fetuses, fetal organs (Fig.1F), and placentae (Fig.1G) and supporting volumetric segmentation and analysis (Fig.1H).

**Figure 1.**
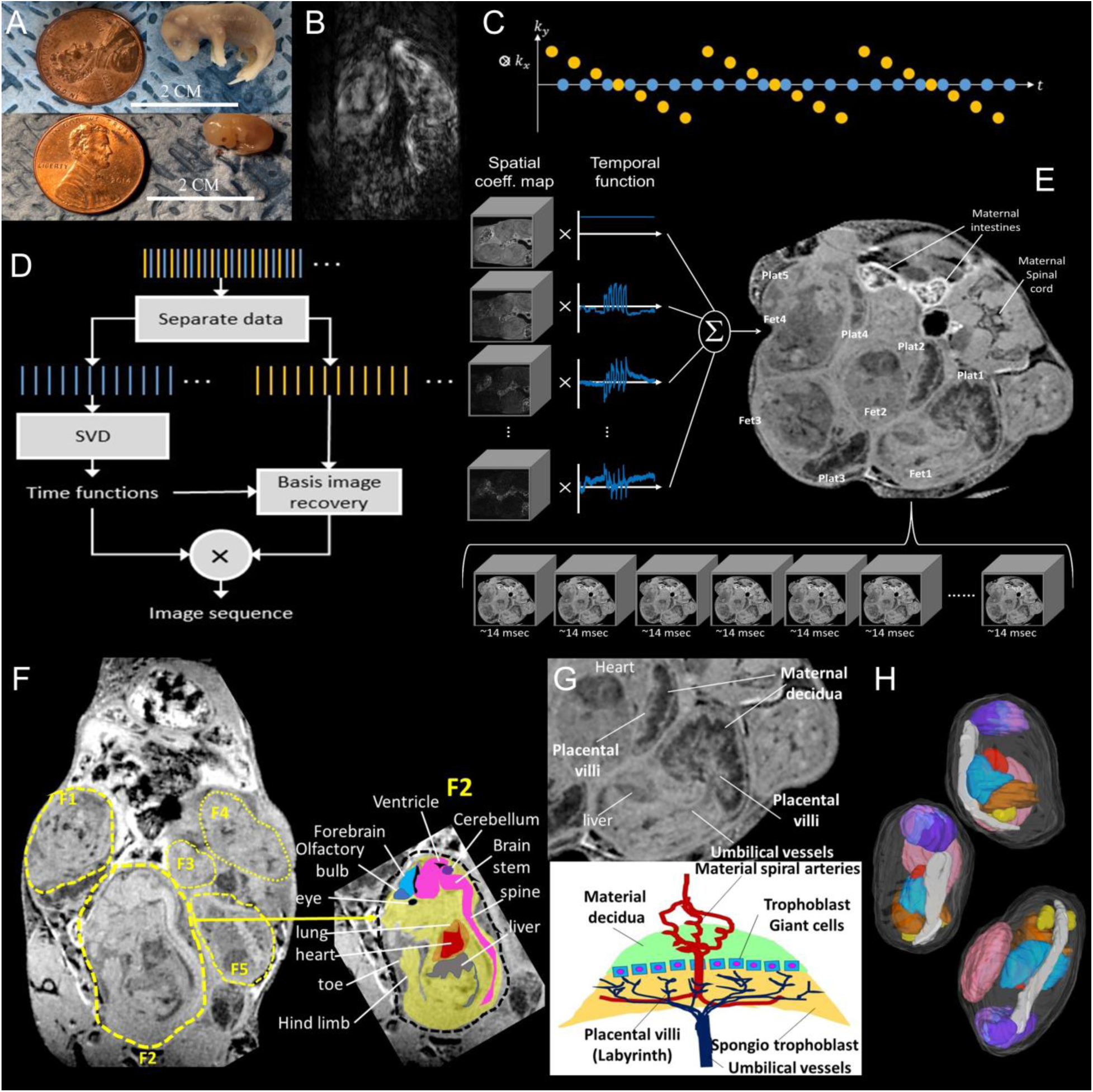
Low-rank 4D *in utero* MRI (4D-uMRI) with sub-Nyquist scheme. (A) Mouse fetuses on the embryonic day E16.5 (top) and E14.5 (bottom) compared to a penny. The scale bar indicates 2 cm. (B) A conventional fast MRI scheme with echo planar imaging (EPI) of a pregnant female mouse suffer from motion artifacts from multiple fetal heart beats, fetal motions, and maternal respiration. (C) Illustration of *k*-t sampling patterns. 4D-uMRI splits imaging sequences into 2 parts: one spatial basis (orange) and one temporal (blue) functions. Acquisition alternates between two sets of data: basis imaging data (orange) with high spatial and low temporal resolution, with full k-space; and training data (blue) with high temporal resolution with limited k-space coverage for fast acquisition. (D) Flowchart of the image reconstruction algorithm. The acquisition data are separated to two parts, the imaging basis data (orange) and temporal basis data (blue). The interleaved training data (blue) are used to determine the temporal basis function with the singular value decomposition (SVD). Then the basis image functions are determined by fitting the temporal basis to the remainder of the imaging data, to generate 4D time series. (E) Illustration of combining D1 temporal function and D2 basis function to generate full time series with reduced *k*-t steps to achieve accelerated low-rank imaging. (F) One time frame of the 4D MRI of a pregnant female mouse showing 5 fetuses (F1 to F5) on embryonic day E16.5 with no motion artifact. Each time frame of the 4D-uMRI is a 3D isotropic image stacks, thus it can be re-oriented for any viewing angle. This angle showing the sagittal view of fetus F2. The colored drawings show corresponding brain and visceral organ segmentation. The 4D-uMRI was acquired at Bruker 7-Tesla 70/30 USR scanner with a 35-mm quadrature volume coil, 120 μm x 120 μm x 120 μm isotropic resolution, 10° flip angle (FA) and total 45-min scan time. (G) One frame from the 4D-uMRI stack showing 2 placentae without motion artifact. The colorful drawing shows mouse placental anatomy. (H) The *in vivo* 4D-uMRI is sufficient for segmentation for volumetric analysis. 3 mouse fetuses are shown with various ROIs segmented: Lavender – brain, Red - heart, Blue - lung, Brown -liver, Yellow - kidney, Gray - spine, Pink - placenta.

Standard fMRI acquires low-resolution 2D BOLD images (Fig 2A, yellow blocks) superimposed on separately acquired high-resolution 3D anatomical images (Fig 2A, purple blocks), requiring additional preparation for plane alignment (Fig 2A, blue block) and risking resolution mismatch and co-registration errors, especially in multi-fetal pregnancy with random orientations (Fig.2A).

**Figure 2.**
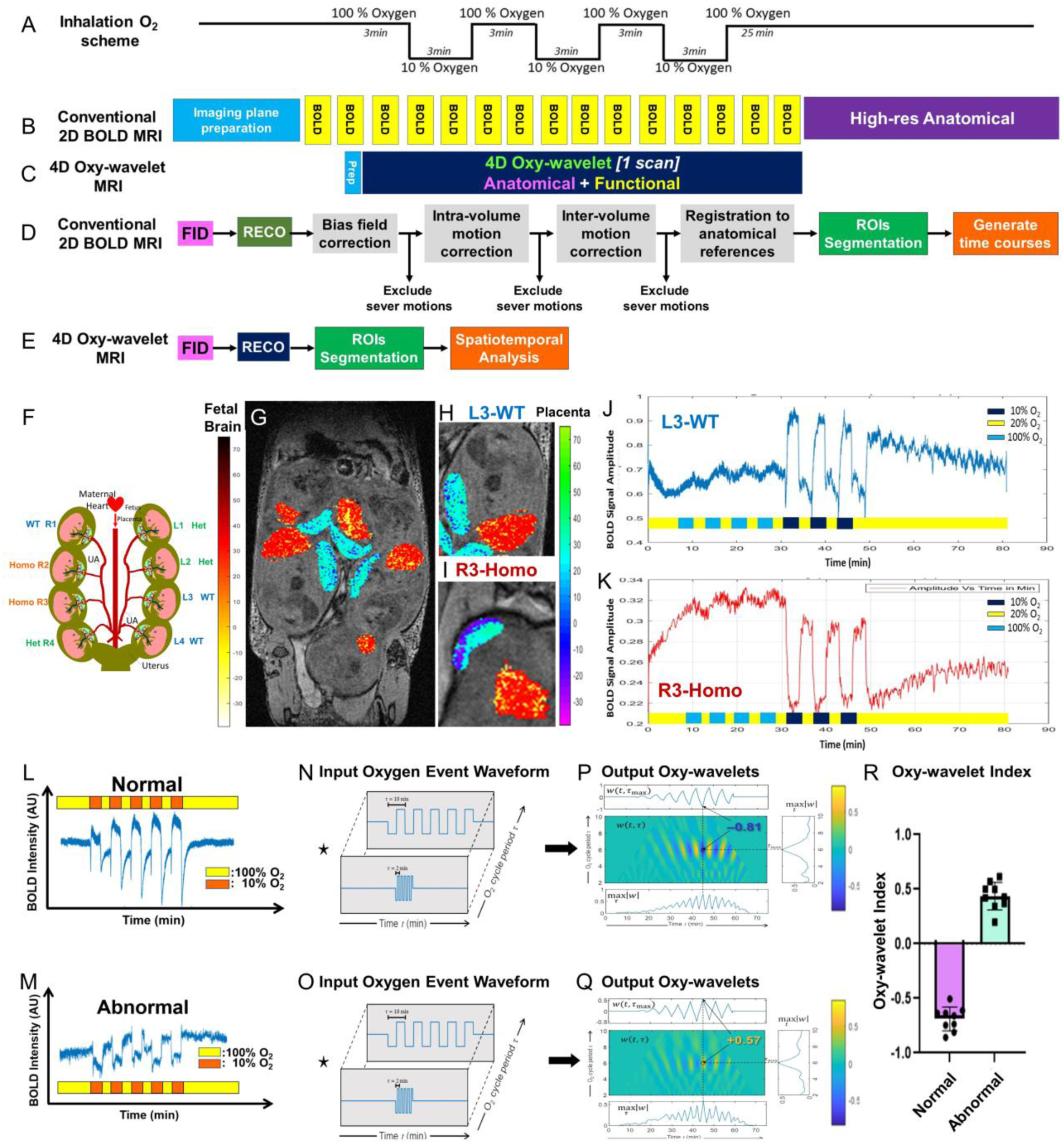
4D Oxy-wavelet MRI scheme and for fetal brain and placental capability to compensate acute hypoxia challenges in *Sap130* fetuses on E16.5. (A-E) 4D Oxy-wavelet MRI experimental protocol and processing pipeline (A) Maternal inhalation O_2_ scheme. (B) Conventional 2D BOLD MRI acquisition scheme with sequential acquisition of each functional parameter with resolution mismatch for functional and anatomical scans. (C) acquisition scheme: integrated single 4D scan for isotropic anatomical and functional scan. (D, E) processing pipeline for (D) Conventional 2D BOLD MRI (E) 4D Oxy-wavelet MRI. FID: free induction decay, Reco: reconstruction. ROI: region of interest. (F-K) 4D Oxy-wavelet MRI of a *Sap130* pregnant female. (F) Drawing of a litter of 8 fetuses on E16.5 with 3 WT (blue R1, L3, L4), 3 heterozygous (*Sap130^+/m^*, green L1, L2, R4) and 2 homozygous (*Sap130^m/m^*, orange R2, R3) mutants. (G-I) Voxel-wise integration of BOLD signals during hypoxia. The red-yellowish color scales represent fetal brain BOLD summation, whereas the blue-greenish color scales represent the placental BOLD summation. (G) the litter in Fig.4F, (H) L3-WT, (I) R3-homozygous mutant with poorer heart and brain outcomes. (J, K) Placental BOLD signals of embryo (J) L3-WT and embryo (K) R3-homozygous). Y-axis: placental BOLD signals. X-axis: time. Yellow bars: normoxia (20% O_2_), light blue bars: hyperoxia (100% O_2_), dark blue bars; hypoxia (10% O_2_) periods. (L-R) Oxy-wavelet waveform analysis. (L, M) Temporal fetal brain BOLD responses with normal (L) and abnormal (M) mitochondrial functions. Yellow periods: high oxygen (100% O_2_); red: hypoxia (10% O_2_). (N, O) Cycling input oxygen event waveforms with various frequency. (P, Q) Output feature Oxy-wavelets for a normal (P) and abnormal (Q) fetal brains. The location and sign of the peak value reveal the oxygen response timing and direction, respectively. The negative peak indicates an opposite BOLD response (brain sparing: higher tissue oxygenation when external oxygen decreases), whereas the positive peak indicates correlated BOLD response (lower tissue oxygenation when external oxygen decreases), distinguishing naïve and irradiated responses. (R) Oxy-wavelet Index for normal (purple, negative) and abnormal (green, positive) mitochondrial functions.

Our method acquires one single 4D scan (Fig.2B, dark blue block) covering the entire volume, with minimal preparation time (Fig 2B, light blue block). Furthermore, conventional fetal imaging requires bias-field correction^44–46^, intra-volume and inter-volume motion corrections, and co-registration to the anatomical reference (Fig. 2C) before regions of interest (ROIs) can be segmented. On the contrary, our method (Fig.2D) is motion-resolved so no motion correction or exclusion is needed while the anatomical and functional information shares the same resolution and geometry, so no co-registration is required.

### 4D in *utero* Oxy-wavelet MRI can probe *in vivo* mitochondrial dysfunction in fetal brains and placentae

4D Oxy-wavelet MRI leverages oscillating hypoxia challenges (Fig.2E) to probe *in vivo* mitochondrial functions. To maintain oxygen homeostasis^47,48^, acute hypoxia triggers fast cardiorespiratory reflexes with hyperventilation and sympathetic responses to increase pulmonary gas exchange and blood flow to bring more oxygen to vital organs such as brains and hearts. <u>Systemic hypoxia responses</u>^47,49^ are initiated by oxygen-sensing arterial chemoreceptors^47,49^ in the carotid body, pulmonary arteries, ductus arteriosus, adrenal medulla, and lung neuroepithelial bodies. Oxygen sensing of these systems requires redox signaling of mitochondrial ETC complex I^49^ and, <u>at the cellular level</u>, oxygen sensing requires mitochondrial ETC activity^49–53^ and ROS-mediated signaling^50,52,54^ for transient compensatory activation of respiratory chain to maintain oxygen homeostasis and metabolism. Mitochondrial dysfunction therefore impairs oxygen sensing and tissue oxygenation, and4D Oxy-wavelet MRI leverages this as a biomarker of mitochondrial dysfunction in the brain and placenta.

Efficient fetal oxygen delivery, regulated by placental-fetal brain interplay, is crucial for fetal brain development. *SAP130* encodes a Sin3A-associated protein 130^55^, a chromatin modifying protein in a Sin3A-histone deacetylase (HDAC) repressor complex^56,57^ that functions as a transcriptional repressor of various fetal development genes, and regulating mitochondrial energy productions^58^. Crossing heterozygous breeding pairs generates fetuses with different genotypes in the same maternal environment to distinguish maternal influences from intrinsic fetal factors. In one litter of eight fetuses (Fig.2F), three were WT (blue R1, L3, L4), three were heterozygous (*Sap130^+/m^*, *^green^* L1, L2, R4) and two were homozygous (*Sap130^m/m^*, orange R2, R3). When maternal inhaled oxygen switched normoxia (20% O_2_, Fig.2 J, K, yellow periods) to hyperoxia (100% O_2_, Fig.2 J, K, light blue periods), both L3 WT placenta (Fig.2J) and R3 homozygous placenta (Fig.2J) increased BOLD amplitudes. On the other hand, when subjected to acute oscillating hypoxia stress (10% O_2,_ Fig.2 J, K, orange-redish periods), a L3 WT placenta (Fig.2J) was able to compensate for the hypoxia and adjusted to an even higher oxygenation state (10% O_2_, Fig.2 J orange-redish periods) after the initial transient dip. On the contrary, R3 homozygous placenta (Fig.2K) was unable to compensate, with placental BOLD signals remaining low during the hypoxia challenges (Fig.2K orange-redish) until the mother’s inhalation returned to normal. Fetal brain BOLD responses parallel those of placentae.

Voxel-wise integration of BOLD levels during hypoxia revealed that L3-WT fetuses had larger placentae and brains with high BOLD levels, whereas R3-homozygous fetuses had smaller placentae and brains with predominantly low BOLD voxels near the decidua (Fig.2G–I).

During oscillating hypoxia (10% O_2_) challenges (Fig.2 L, M, orange-red periods), normal fetal brains (Fig.2L) could overcome hypoxia to quickly adjust to a higher oxygenation state after the initial transient BOLD signal drop. Conversely, abnormal fetal brains (Fig.2M) showed passive response to hypoxia, with BOLD levels staying low.

We tested the capability of 4D Oxy-wavelet MRI to probe *in vivo* mitochondrial dysfunction in *Ndufs4* fetal brains with mitochondrial ETC complex I defects (Fig.3 A-F) and in placentae of peri-conceptional alcohol exposure (Fig.3 G-I).

**Figure 3.**
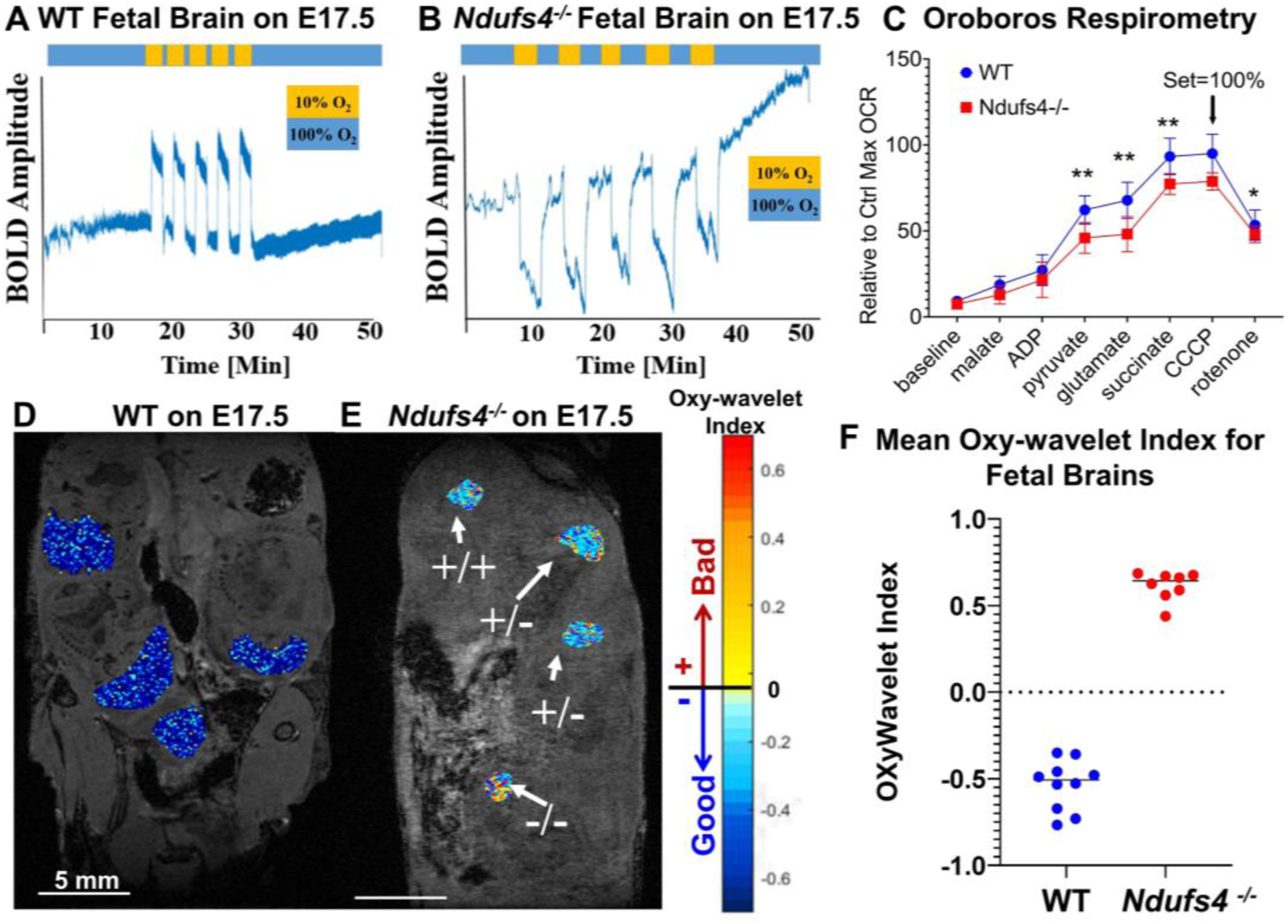
*In utero* 4D Oxy-wavelet MRI for genetic mutant *Ndufs4* fetal brains. (A-F) *Ndufs4* encodes a sub unit in the mitochondrial ETC complex I. Homozygous *Ndufs4^−/−^* mutants have mitochondrial ETC dysfunction. (A, B) Temporal BOLD responses of fetal brains on E17.5 for WT (A) or homozygous *Ndufs4^−/−^* (B). Y-axis: BOLD amplitude of fetal brains. X-axis: time (minutes). Blue periods: high oxygen (100% O_2_); orange: hypoxia (10% O_2_). (C) *Oroboros* mitochondrial respirometry of fresh mitochondria isolated from fetal brains on E17.5. Y-axis: oxygen consumption. X-axis: time. Activity levels were normalized with mitochondria protein amount. Substrates and inhibitors were added stepwise: cytochrome-C (CytC), malate (Mal), adenosine diphosphate (ADP), pyruvate (Pyr), glutamate (Glu), succinate (Suc), rotenone (Rot), carbonylcyanide-3-chlorophenylhydrazone (CCCP). The oxygen consumption rates were normalized to the highest rate after adding CCCP. Blue – WT (n=3), Red - *Ndufs4^−/−^* (n=3). (D, E) Voxel-wise Oxy-wavelet index maps on E17.5 for WT (D) and *Ndufs4^−/−^* (E) litters. Positive Oxy-wavelet indexes – bad; Negative Oxy-wavelet indexes – good. (F) Mean Oxy-wavelet indexes for fetal brains. Blue – WT (n=10), Red - *Ndufs4^−/−^*(n=8).

Mutations in *NDUFS4* (5q11.2), encoding a complex I subunit, often cause Leigh Syndrome (LS), a severe childhood-onset neurodegenerative disease affecting ∼1 in 40,000 births^59–62^. In *Ndufs4^tm1.1Rpa^*/J LS mice, exon 2 of *Ndufs4* is excised, and homozygous mice develop progressive, lethal neurodegeneration with death before 8 weeks^62,63^. 4D Oxy-wavelet MRI was used to detect homozygous *Ndufs4* fetal brains at E17.5 in a blinded manner (Fig.3A–F).Under oscillating hypoxia challenges (Fig.3 A, B, orange periods), WT fetal brains actively compensated, maintaining higher oxygenation state (Fig. 3A). Most WT fetal brain voxels (Fig.3D) showed good/negative (Fig.3D blueish color scales) Oxy-wavelet indexes. *Ndufs4* KO fetal brains with intrinsic complex I failed to compensate to hypoxia (Fig.3B orange periods), showing persistently low BOLD levels during hypoxia until the mother’s inhalation switching back to normal. *Ndufs4-/-* fetal brains (Fig.3 E, -/-) displayed positive/bad (Fig.3E yellow/orange/red-ish color scales) Oxy-wavelet indexes, with very few voxels being normal. Median Oxy-wavelet indexes across litters were negative in WT fetal brains (Fig. 3F blue) (Fig.3F blue), whereas all *Ndufs4* KO fetal brains showed positive/bad Oxy-wavelet indexes (Fig.3F red). Mitochondrial Oroboros respirometry (Fig.3 C) confirmed reduced oxygen consumption rate (OCR) and compromised complex 1 function in *Ndufs4-/-* brains (Fig.3 C red) compared to the WT controls (Fig.3C blue).

### 4D Oxy-wavelet MRI can probe *in vivo* mitochondrial dysfunction in adult brains in a spatially specific manner

4D Oxy-wavelet MRI was expanded to adult rodent brains to detect mitochondrial dysfunction in a brain-region-specific manner (Fig.4A). Reconstructed 3D volumes (Fig.4A.1) were registered to the Allen mouse brain atlas^64–67^ (72 regions) (Fig.4A.2) for brain region parcellation (Fig.4A.3 Region-specificc BOLD signal profiles (Fig.4A.4) were extracted for Oxy-wavelet analysis (Fig.4A.5) to generate whole-brain 3D Oxy-wavelet index maps (Fig.4A.6).

**Figure 4.**
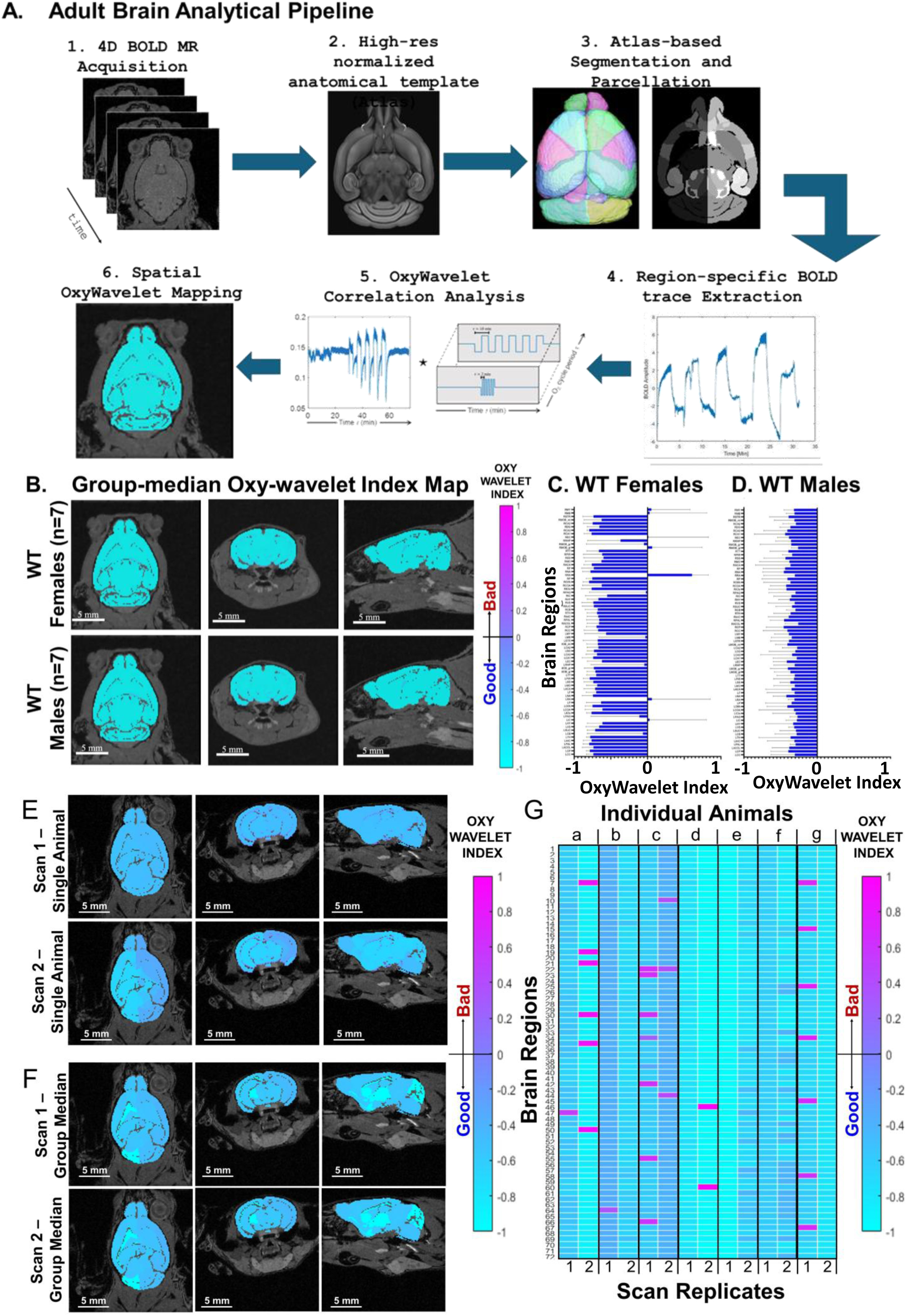
4D Oxy-wavelet MRI in adult brains. (A) 4D Oxy-wavelet MRI analysis pipeline for adult brains. <1> 4D Oxy-wavelet MRI acquisition, <2> Normalized high-resolution anatomical template, <3> atlas-based segmentation and brain region parcellation, <4> brain region-specific temporal BOLD signal profiles, <5Ϭ Time-frequency Oxy-wavelet waveform analysis, <6> Output Oxy-wavelet maps. (B) Group median 3D Oxy-wavelet index maps for WT (top) females (n=7) and (bottom) males (n=7). (C) Group median Oxy-wavelet indexes for WT females (n=7) for each of the 72 brain regions. Error bars – standard deviation. (D) Group median Oxy-wavelet indexes for WT males (n=7) for each of the 72 brain regions. Error bars – standard deviation. (E) 3D Oxy-wavelet index maps for 2 repeated scans of the same animal. (top) scan 1, (bottom) scan 2 of the same animal. (F) Group median of 7 animals with repeated scans. (top) scan 1, (bottom) scan 2. (G) Individual Oxy-wavelet indexes of each brain region for 7 animals (a-g), with 2 scans (scan 1 and 2) for each animal.

Group-median Oxy-wavelet maps for WT mice (Fig.4C) showed largely negative/good Oxy-wavelet indexes (Fig. 4D blue-ish color scales), indicating intact ETC functions. Region-wise analysis found that WT males (Fig.4D) showed negative/good indexes in all 72-brain regions, whereas females (Fig.4C) had positive/bad indexes in left inferior colliculus, left reticular nucleus, right reticular nucleus, right olfactory bulb glomerular layer, right entorhinal cortex, right midbrain and right medulla.

### 4D Oxy-wavelet MRI in Normal Adult Brains – Reproducibility and Specificity

To assess reproducibility, seven WT mice underwent 4D Oxy-wavelet MRI on 2 days (Fig.4 E, F). Oxy-wavelet index maps were similar within individual animals (Fig.4 E, F) and in group-averaged maps (Fig.4F) across days. Among 504 regional comparisons (Fig. 4G), 479 had the same index sign, yielding 95% reproducibility, and 27 of 1008 regions exhibited positive (bad) indexes across 14 scans, corresponding to a 2.7% false positive rate (FPR).

Specificity for mitochondrial function versus hemodynamics was tested by increasing CBF with adenosine (hyperemia) and by selectively inhibiting mitochondrial complex I with rotenone. Vasodilator adenosine injected during oscillating oxygen flipping (Fig. 5A) did not change temporal BOLD waveforms (Fig. 5B) or Oxy-wavelet maps (Fig. 5C). There were no changes in Oxy-wavelet indexes for each brain region (Fig.5D) before (Fig.5D blue) and after (Fig.5D red) adenosine administration.

**Figure 5.**
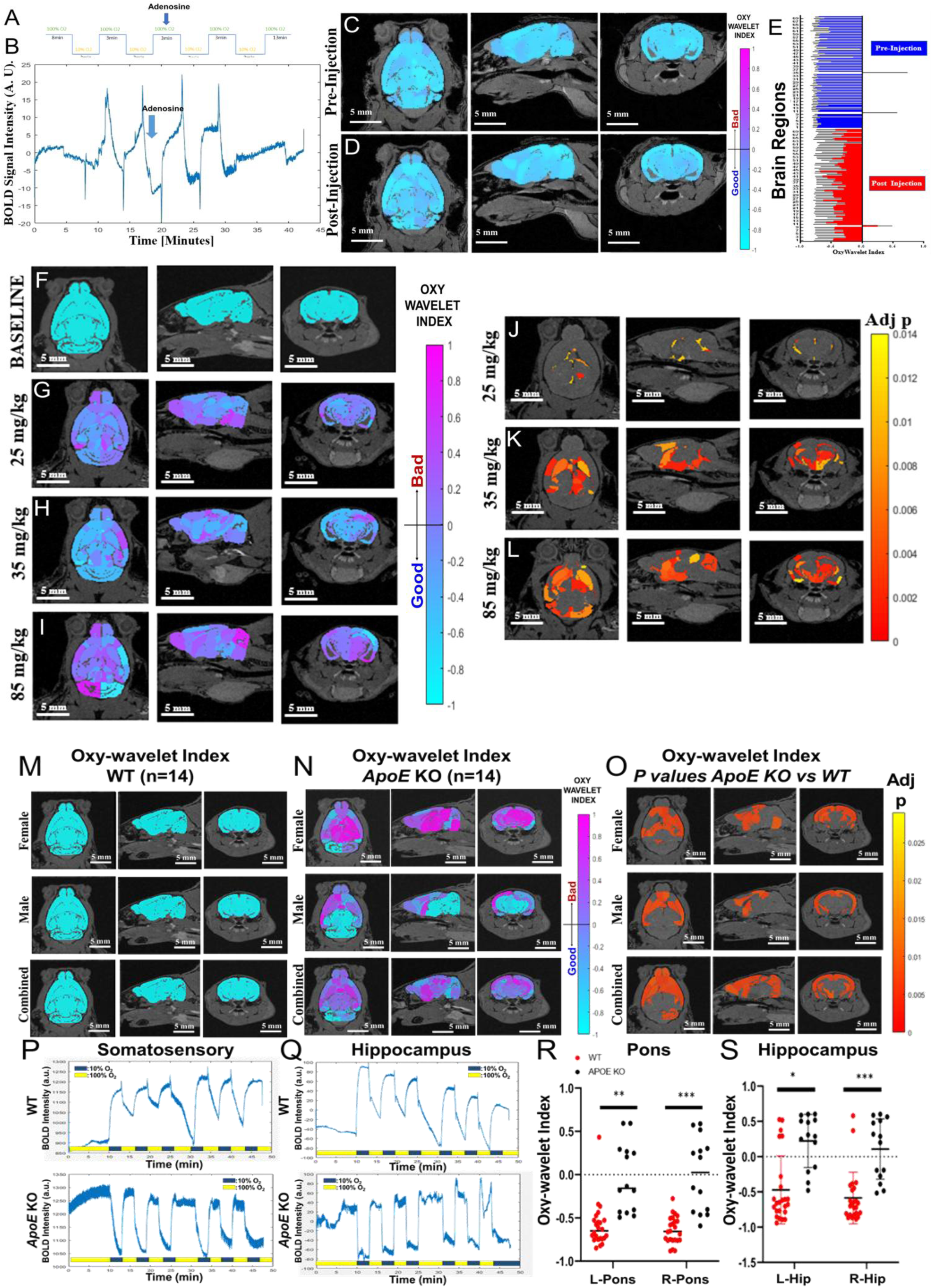
*In vivo* 4D Oxy-wavelet index mapping for adult mouse brains with adenosine, rotenone or ApoE-/- mice. (A) Input oxygen-flipping scheme with adenosine administration in the middle. (B) Temporal brain BOLD profiles with Adenosine administration between the 2^nd^ and the 3^rd^ hypoxia challenge blocks. (C,D) Group median Oxy-wavelet index mapping in WT mice before (C) and after (D) Adenosine (250 mg/kg/min, n=11) administration. (E) Group median Oxy-wavelet indexes for each brain region before (blue) and after (red) Adenosine injection. (F-I) Group-median Oxy-wavelet index maps for baseline (F) or rotenone (G-I) with accumulated doses of 25 mg/kg (G), 35 mg/kg (H), or 85 mg/kg (I). (J-L) Adjusted p value maps comparing rotenone-treated (G-I) to baseline (F) controls with accumulated doses of 25 mg/kg (J), 35 mg/kg (K), or 85 mg/kg (L). (M) Group median Oxy-wavelet index mapping in WT females (top, n=7), WT males (middle, n=8) or combined WT males and females (bottom, n=14). (N) Group median Oxy-wavelet index mapping in *ApoE* KO females (top, n=7), *ApoE* KO males (middle, n=8) or combined *ApoE* KO males and females (bottom, n=14). (O) Adjusted p value maps of comparing *ApoE* KO vs WT. females (top, n=7), males (middle, n=8) or combined males and females (bottom, n=14). (P, Q) Temporal BOLD signal profiles in somatosensory cortex (P) or hippocampus (Q) for a WT (top) or *ApoE* KO (bottom) mice. (R, S) Oxy-wavelet indexes of individual animals at pons (R) or hippocampus (s). Red: WT (n=14), Black: *ApoE* KO (n=14). ** *p*< 0.01, *** *p*<0.001.

In contrast, rotenone^68–70^ treated animals showed massively compromised Oxy-wavelet indexes (Fig.5 G-I purple-ish color scale, bad function) in a dose-specific manner. Mice received 5 mg/kg/week Rotenone injections to reached cumulative doses of 25 mg/kg (Fig.5G), 35 mg/kg (Fig.5H), or 85 mg/kg (Fig.5I). Compared with sham controls, adjusted *p*-value maps revealed dose-dependent regional changes: few significant regions at 25 mg/kg (Fig. 5J), more extensive involvement at 35 mg/kg (Fig 5K) including thalamus, striatum, primary motor cortex, and piriform cortex, and widespread changes at 85 mg/kg (Fig. 5L) involving thalamus, striatum, primary motor cortex, piriform cortex, and cerebellum.

### 4D Oxy-wavelet MRI can detect mitochondrial dysfunction in ApoE KO adult brains

Apolipoprotein E (*APOE*)^71–75^ isoform is the strongest genetic risk factor for late-onset Alzheimer disease (LOAD)^76–78^. Mitochondrial dysfunction^18,24,79,80^ is well recognized in LOAD. In *ApoE*-/- mice, 4D Oxy-wavelet MRI revealed compromised indices, (Fig.5M-S, positive/bad function, purple-ish color scales) with females (Fig.5N Top) being worse than males (Fig.5N middle). *P* value maps (Fig.5O) comparing *ApoE-/-*(Fig.5N) vs WT (Fig.5M) found differences in the somatosensory cortex, hippocampus, amygdala, and pons (Fig.5O). Temporal BOLD waveforms from somatosensory cortex (Fig.5P) and hippocampus (Fig.5Q) showed distinct waveform characteristics between WT (Fig. 5P, Q, top) and *ApoE*-/- (Fig. 5P, Q bottom). WT brains actively compensated for oscillating hypoxia (Fig. 5P, Q dark blue periods) with BOLD levels above baseline, whereas *ApoE*-/- brains failed to compensate (Fig. 5P, Q dark blue periods) with BOLD levels staying lower than the baseline. Most WT brains (Fig. 5R, S red) displayed negative/good Oxy-wavelet indexes but more *ApoE* KO mice (Fig. 5R, S black) showed positive/bad Oxy-wavelet indexes in pons (Fig.5R) and hippocampus (Fig.5S). 4D Oxy-wavelet MRI thus detects intrinsic mitochondrial dysfunction caused by *ApoE* deletion.

### 4D Oxy-wavelet MRI can be safely conducted in humans without hypoxia

4D Oxy-wavelet MRI was safely implemented in humans using intermittent 30-sec hyperventilation (HV) to trigger mitochondrial ETC responses. In a single, healthy human volunteer without MRCD history (Fig. 6), 5 min 4D scan using a multichannel head coil with normal breathing (Fig. 6A, E) or HV (Fig. 6, F-I) provided both functional BOLD and anatomical information (voxel size 1 × 1 × 4 mm) without separate fMRI–anatomy registration, with SNR ∼40.4.

**Figure 6.**
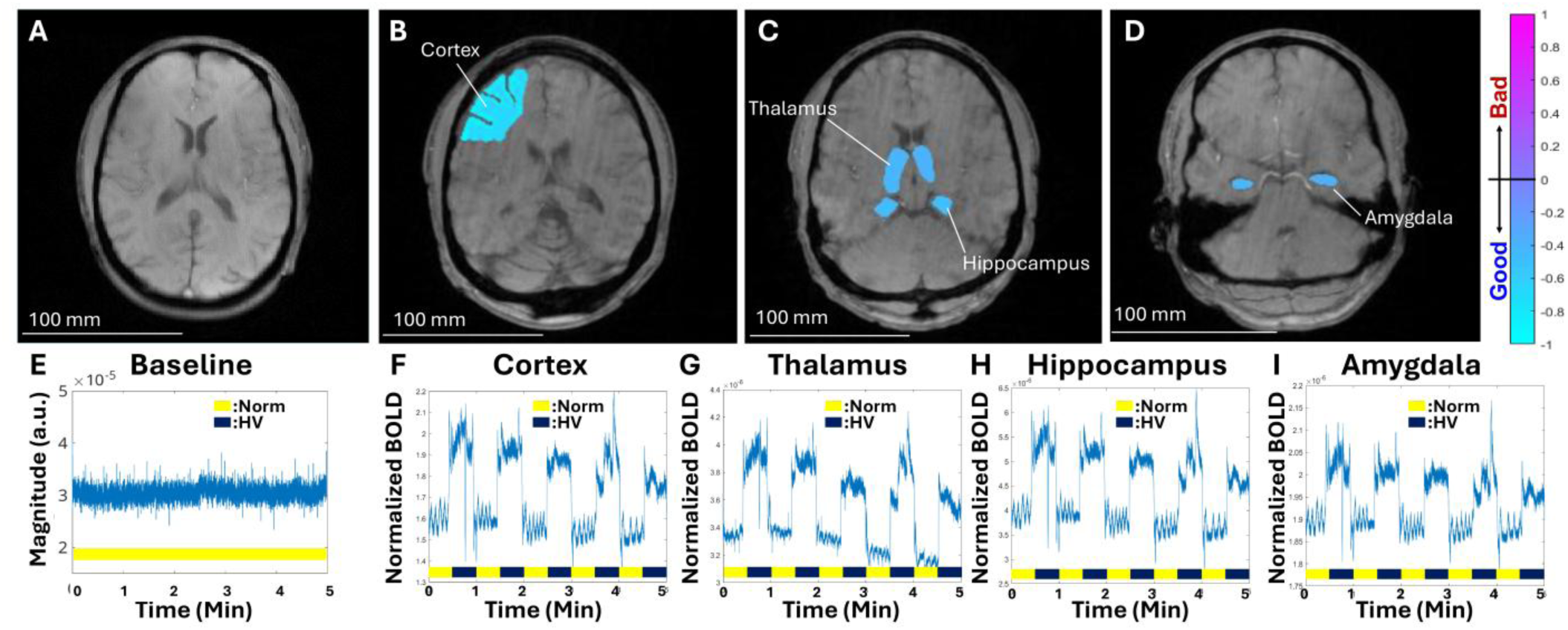
Human 4D Oxy-wavelet MRI of a healthy 38-year-old Caucasian male with no history of MRCD. (A) 4D MRI with normal breathing. (B-D) Regional Oxy-wavelet index maps, negative Oxy-wavelet index - good, blueish color; positive Oxy-wavelet index -bad, purple-ish color for (B) Cortex, (C) Thalamus and Hippocampus, (D) Amygdala. (E) Brain signal with normal breathing. (F-I) Regional Oxy-wavelet signals with intermittent 30-sec hyperventilation (HV) for (F) Cortex, (G) Thalamus, (H) Hippocampus, (I) Amygdala. Yellow periods: “Norm” normal breathing; dark blue periods: “HV” hyperventilation. 4D scans were acquired at a 3-Tesla Siemens Skyra scanner via a 20-channel head coil with the following parameters: echo time TE: 7.5 ms, repetition time TR: 10 ms, flip angle FA = 8^0^, total scan time: 5 min.

Wavelet analysis in cortex (Fig. 6B, F), thalamus (Fig. 6C, G), hippocampus (Fig. 6C, H), and amygdala (Fig. 6D, I) showed negative (“good”) Oxy-wavelet indexes (Fig. 6B-D, blueish color scales) and favorable waveforms (Fig. 6F-I). During HV, BOLD (Fig. 6F-I, dark blue periods) levels rose above resting baseline, indicating normal ETC function, and the temporal resolution (∼20 ms) allowed resolution of respiratory fluctuations during normal breathing. These data demonstrate that 4D Oxy-wavelet MRI can be safely performed in humans with HV and is readily translatable.

## Discussion

4D Oxy-wavelet MRI simultaneously probes *in vivo* mitochondrial function in fetal brains, placentae, and adult brains across the lifespan with spatial specificity. A special low-rank *k*-t space-sampling scheme with partially separated temporal and spatial functions achieves high spatial-temporal resolution 4D fMRI with no motion artifacts. Physiological BOLD responses are captured in the same scan as anatomy, enabling fMRI at anatomical imaging resolution and SNR and avoiding conventional fMRI limitations. By exploiting the requirement for ETC activation to maintain oxygen homeostasis during hypoxia or HV, 4D Oxy-wavelet MRI detects mitochondrial dysfunction in fetal brains and placentae with genetic mutations (*Sap130, Ndufs4*), or fetal injury. Generating multiple fetal genotypes within a shared maternal environment enables dissection of fetal versus maternal factors along the placenta–brain axis and supports differential diagnosis of individual placentae and fetal brains within the same litter, with validation from mitochondrial Oroboros respirometry.

In adult brains, atlas-based segmentation allows region-specific evaluation of mitochondrial dysfunction and 3D Oxy-wavelet index mapping across the brain. 4D Oxy-wavelet MRI detects brain-region-specific mitochondrial abnormalities due to genetic mutations (*ApoE*) or pharmacological insults (rotenone), including hippocampal and somatosensory cortex deficits consistent with known *ApoE*-/- phenotypes^81–83^. The method also reveals sex differences in a LOAD mouse model, with females showing more dysfunctional regions than males, and distinguishes mitochondrial impairment from isolated hemodynamic changes, as adenosine-induced hyperemia does not alter Oxy-wavelet readouts, whereas chronic rotenone exposure induces dose-dependent regional dysfunction consistent with striatal vulnerability in Parkinsonian rodent models^70,84–86^.

4D Oxy-wavelet MRI uses time-frequency waveform cross-correlation to generate wavelet space to evaluate good or bad functions. Consequently, 4D Oxy-wavelet MRI has excellent reproducibility compared to conventional fMRI studies. Specifically, we demonstrate 5x to 25x better FPR than conventional fMRI (10-70%)^87–91^. Conventional fMRI relies only on BOLD signal amplitudes to detect resting state or tasks induced BOLD signal changes. Typical BOLD signal changes are small (≤4%) so extensive statistical methodology is necessary, which can inflate FPR^87–91^. 4D Oxy-wavelet MRI is robust with large signal changes (∼30%) so no statistical maneuvers are needed to evaluate BOLD responses.

The method implements non-rigid input oxygen waveforms with varied frequencies to accommodate imperfect experimental timing, as oxygen is manually switched by altering O_2_ and N_2_ flows supplied via a nose cone. Wavelet analysis with variable frequencies, durations, and amplitudes supports robust readouts under realistic conditions, including irregular square-wave inputs and transient BOLD drops during initial oxygen sensing. This approach can also accommodate variability in human HV performance and enables automated detection of passive versus active brain responses, providing a basis for future AI-based diagnostics.

4D Oxy-wavelet MRI uses a smaller flip angle (10°) than conventional BOLD sequences (∼80°) because of shorter TE/TR (2.3/8.7 ms vs 32/2280 ms), yielding a lower Ernst angle and faster acquisition.

Initial development on preclinical scanners provides a foundation for human translation. SNR scales with field strength, voxel volume, and √ (scan time). In mice at 7T, 4D Oxy-wavelet MRI used 120-μm and 78-μm isotropic resolution for fetal and adult brains. Human fetal and adult imaging requires ∼1.0 mm resolution, providing >500× SNR gain that is tempered by lower clinical field strengths (1.5–3 T), yielding an overall estimated SNR boost of (1.5T/7T)*(1.0 mm/120 μm)^3^ ≈ 124, not accounting for multichannel coil benefits. Equivalent SNR to 64 min mouse scan is thus estimated to be achievable in 64min/√124 = 5.7 min.

Even without breathing maneuvers, the low-rank 4D MRI can acquire a full 3D volume without motion artifact in 5 minutes. It does not require prescriptions of specific imaging planes, thus needs very little preparation time. This can allow fetal or neonatal imaging without anesthesia. Anesthesia is a common practice for infant or young pediatric neuroimaging to avoid motion artifacts. Anesthesia imposes increased adverse risks for patients with mitochondrial disease. The low-rank 4D MRI can facilitate anatomical MRI in infants or young children without anesthesia. This can potentially allow detecting *in vivo* mitochondrial dysfunctions in patients for individualized diagnosis, prognosis, and physiology-based surrogate endpoints for clinical trials.

## Data Availability Statement

All data pertinent to this work will be made available via a public data repository hosted on the Neuroimaging Tools & Resource Collaboration online page (NITRC).

## Author Contributions

The author contributions are as follows: DREC— Idea conception, data processing, formal analysis, writing original manuscript, review and edit manuscript. SH—Data acquisition. TBS—Data acquisition. KES—Data acquisition. CR—data processing. SM—data processing. NWC—Animal handling, data analysis. DW—animal handling, data analysis. MCS—animal handling, data analysis. SW—data acquisition. CWL—animal model, review and editing. SB—data analysis. ESG—manuscript review and editing. AGC—project conception, data analysis, formal analysis, manuscript writing, review and editing. YLW—project conception, project administration, formal analysis, data acquisition, manuscript review, writing, and editing.

## Declaration of Competing Interests

The authors declare they have no competing interests to disclose in regards to the work contained within this submission.

## Acknowledgements / Funding

This study was supported in part by funding to YLW from NIH (EB023507, NS121706, NS142555), AHA (18CDA34140024), and DoD (W81XWH1810070, W81XWH-22-1-0221).

## Methods

### Ethics Statement and Animal Handling

Mice were housed and cared for according to the animal protocol (IACUC protocol # 22050950 and 25066690) approved by the Institutional Animal Care and Use Committee, University of Pittsburgh and the Animal Care and Use Review Office (ACURO), US Army Medical Research and Development Command (USAMRDC). Animals were housed in cages with centralized filtered air/clean water supply system in a secure AALAS certified vivarium in the Rangos Research Center of the Children’s Hospital of Pittsburgh of UPMC. Animal care is provided seven days a week based on the NIH Guide for *the Guide for the Care and Use of Laboratory Animals.* Animals were kept in a 12:12 hour dark/light cycle with *ad libitum* food and water.

All data was generated and analyzed in accordance with ARRIVE^114^ guidelines, including protocols for blinding, randomization, counterbalancing, inclusion of proper controls, and appropriate statistical power.

Breeding pairs of WT C57BL/6J (Jax Stock #000664), *Ndufs4^tm1.1Rpa^*/J (Jax Stock #027058), and *ApoE^tm1Unc/^J* KO (Jax Stock #002052) were obtained from Jackson Laboratories then bred in house. *Sap130* mice were generated in house via mutagenesis screening^115^.

### Time-resolved free-breathing/gating-free 4D *in utero* Oxy-wavelet MRI

We have successfully implemented a novel 4D time-resolved *in utero* MRI (4D-uMRI) method that can image each fetus simultaneously with no motion artifacts and can capture functions at the same time. The two key features of this novel fetal MRI approach are (1) the ability to express incoherent fetal and maternal motion with reduced degrees of freedom; and (2) a sparse sampling scheme to accelerate acquisition and increase temporal resolution. This methodology allows assessments of anatomical structures and observe hemodynamic function and its relationship to the developing fetal brain. Multiple fetuses are imaged without motion artifact. Fetal brain, heart, liver, and placental structure are identified.

We use a hybrid low-rank and sub-Nyquist sparse sampling technique^92–99^ to perform 4D imaging *in utero*. The low-rank model expresses the image *ρ*(𝐫, *t*) (for voxel location 𝐫 and time *t*) as:

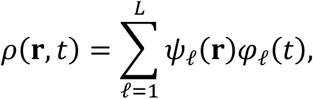

where 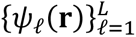 are *L* spatial coefficient maps and where 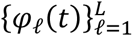 are *L* temporal basis functions. In matrix form, this becomes **x** = 𝚿𝚽, where *X*_*ij*_ = *ρ*(𝐫_***i***_, *t*_*j*_), 𝛹_*ij*_ = *ψ*_*j*_(𝐫_*i*_), and ***Φ***_*ij*_ = *φ*_*i*_(*t*_*j*_). We additionally modeled *ρ*(𝐫, *t*) as being a transform sparse in the wavelet-spectral domain (i.e., we model ℱ_*t*_{*W*_***r***_{*ρ*(𝐫, *t*)}} as sparse, where *W*_***r***_is a spatial wavelet transform and ℱ_*t*_ is the temporal Fourier transform). This approach exploits both image correlation and transform sparsity to allow high spatiotemporal resolution imaging.

A useful data acquisition strategy (Fig.1C) alternates between collecting two sets of data: D1 (Fig.1C, blue), which contains auxiliary “navigator” data collected at a high temporal sampling rate but a limited number of **k**-space trajectories, and D2 (Fig.1C, orange), which contains sparsely sampled (**k**, *t*)-space data with extended **k**-space coverage. This strategy capitalizes on the PS model’s decoupled resolution requirements: the high-speed data in D1 inform the temporal basis functions (i.e., the *φ*’s), and the full–**k**-space data in D2 inform the spatial coefficient maps (i.e., the *ψ*’s), resulting in images with the high temporal resolution of D1 and the high spatial resolution of D2.

The data in Fig. D1 was acquired by collecting every other *T*_R_ (i.e., every other readout, for a final frame rate of 1/[2*T*_R_]), using a center **k**-space trajectory. We then performed image reconstruction (Fig. 1D) in two steps: first by determining the temporal basis 𝚽 from the singular value decomposition (SVD) of the interleaved training data, and then secondly determining the spatial coefficient maps 𝚿 by fitting the temporal basis to the remainder of the imaging data. This second step is calculated according to

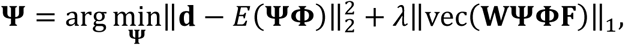

where **d** are the measured data, *E* is the encoding operator comprising spatial encoding and undersampling, 𝐖 is a spatial wavelet transform, **F** is the temporal Fourier transform, and vec(⋅) vectorizes the wavelet-spectral image before calculating the vector 1-norm to promote sparsity.

By combining the D1 temporal function and the D2 basis function, and by exploiting the fact that each frame in the fetal MRI time series has a high degree of spatiotemporal correlation (i.e., they reside in a low-dimensional subspace) to reduce the degrees of freedom required to represent a sequence of images, we achieved rapid imaging speeds beyond that which can be achieved by parallel imaging or fast scanning. The integration of both low-rank imaging^43,100–103^ and compressed sensing^104–107^ performs complementary de-noising, exploiting correlation of images over time and exploits the transform sparsity of the image series.

4D Oxy-wavelet MRI is when applying oscillating hypoxia (Fig.2A) or intermittent hyperventilation (Fig. 6) to trigger mitochondrial ETC responses to maintain oxygen homeostasis to probe mitochondrial functions.

### OxyWavelet Continuous Wavelet Transform

The 4D Oxy-wavelet MRI uses time-frequency analysis of BOLD signals in response to the oscillating hypoxia challenges to probe the integrity of mitochondrial function. Our BOLD signal is event-driven, following the oscillations of the hypoxia challenge; the event waveform resembles a square wave (Fig. 2 N, O). Cross-correlating the BOLD signals (Fig.2 A, B) by template event waveforms (Fig. 2 N, O) at several oscillation frequencies will produce a feature space revealing various details (Fig. 2, P, Q). For example, the location of the peak-magnitude value reveals the midpoint of the oxygen cycling experiment and the oscillation period (neither of which is precisely known *a priori* when oxygen switching is manually triggered). More importantly, the sign of the peak value reveals the direction of oxygen response: a positive peak (Fig. 2Q, bottom) indicates a positive correlation with the oxygen cycling (tissue hypoxia when external oxygen is decreased, a “bad” Oxy-wavelet index), whereas a negative peak (Fig 2R, purple) indicates opposite behavior (brain sparing wherein tissue over-corrects for external oxygen decreases by adjusting to an even higher oxygenation, “good” Oxy-wavelet index).

### Atlas-based brain region segmentation for adult mouse brains

The 3D volume of the reconstructed 4D MRI is first registered to the Allen mouse brain atlas^64,66,67,108^ (72 regions) space as described previously^109^, using ANTx2, a custom MATLAB toolbox and then segmented into gray matter (GM), white matter (WM) and cerebrospinal fluid (CSF) tissue maps using the Unified Segmentation approach^110^ as implemented in statistical parametric maps (SPM)^111^. For the segmentation task, the tissue probability maps (TPMs) are generated based on Hikishima et al.^112^ tissue classification. A weighted image is constructed using the tissue segments of the animal and the TPMs of the template. The weighted images are co-registered using affine and nonlinear B-spline transformation via Elastix package^113^. The resulting parameter for forward transformation is stored to allow a subsequent image transformation from native animal space to rat template space. Parameter files for inverse transformation are also generated and stored to allow a subsequent image transformation from template space to native space (e.g. hemispheric brain mask). Using the files for inverse transformation, we transform the template to the native, 1^st^ volume of the 3D data to create the final brain segmentation mask. Each brain is parcellated into 72 (mouse) without assigning any region of interest (ROI) or region of avoidance (ROA).

### Manual segmentation for fetal brains and placentae

The acquired free-induction decay (FID) was reconstructed with custom algorithm. A single timeframe of the reconstructed 4D time series was arbitrarily selected to generate 3D isotropic imaging stacks. Fetal brains and placentae were manually segmented by blinded observers using the open-source ITK-Snap (www.itksnap.org). The segmented masks were applied to the reconstructed 4D Oxy-wavelet MRI time series for dynamic signal analysis of BOLD summation, Oxy-wavelet index, and oxygen transport time delays.

### Anesthesia for *in vivo* MRI

All mice received general inhalation anesthesia with Isoflurane for *in vivo* imaging. Mice were placed into a clear Plexiglas anesthesia induction box that allows unimpeded visual monitoring of the animals. Induction was achieved by administration of 3% Isoflurane mixed with oxygen for a few minutes. Depth of anesthesia was monitored by toe reflex (extension of limbs, spine positioning) and respiration rate. Once the plane of anesthesia was established, it was maintained with 1-2 % Isoflurane in oxygen via a designated nose cone and the mouse was transferred to the designated animal bed for imaging. Respiration was monitored using a pneumatic sensor placed between the animal bed and the mouse’s abdomen while rectal temperature was measured with a fiber optic sensor and maintained with feedback-controlled warm air source (SA Instruments, Stony Brook, NY, USA).

### In vivo MRI

*In vivo* MRI was carried out on a Bruker BioSpec 70/30 USR spectrometer (Bruker BioSpin MRI, Billerica, MA, USA) operating at 7-Tesla field strength, equipped with an actively shielded gradient system, using a quadrature radio-frequency volume coil with an inner-diameter of 35 mm for transmission and reception.

### *In utero* 4D Oxy-wavelet MRI Acquisition in pregnant mice

4D *in utero* MRI was acquired with the custom pulse sequence with the following parameters: field of view (FOV) = 4.8 cm × 2.8 cm × 2.0 cm, matrix = 355 × 208 × 148, echo time (TE) = 3.475 msec, repetition time (TR) = 7.677 msec, flip angle (FA) = 10°, number of average (NA) = 1, number of repetition (NR) = 12, total scan time = 47 min.

### *In vivo* 4D Oxy-wavelet MRI in adult mice

4D Oxy-wavelet MRI was acquired with the custom pulse sequence with the following parameters: FOV = 2.0 cm × 2.0 cm × 2.0 cm, isotropic resolution (156 μm)^3^, FA =12.5°, TE =2.3 msec, and TR = 9.7 msec, total scan time = 42 min.

### *In vivo* 4D Oxy-wavelet MRI in humans

4D Oxy-wavelet MRI was acquired at a 3-Tesla Siemens Skyra scanner via a 20-channel head coil with the following parameters: TE = 7.5 msec, TR = 10 msec, voxel size = 1 mm × 1 mm × 4 mm, FA = 8^0^, total scan time = 5 min.

### Statistics

Prior to statistical comparison testing, normality testing was performed on each dataset. Normality testing was performed using four different tests and visualization of the dataset QQ-plots. Normality tests performed include: D’Agostino & Pearson test, Anderson-Darling test, Shapiro-Wilk test, and the Kolmogorov-Smirnov test. If any tests failed normality and/or the QQ-plot deviated from linearity, the dataset was assumed to be non-normal in nature. For non-normal datasets, the median ± inter-quartile range (IQR) were used to describe the distribution. Multiple paired or un-paired Wilcoxon signed rank tests with the false discovery rate set to 5% were performed. For normally distributed datasets, unpaired t-tests with unequal variances were performed for comparisons between two groups. For comparisons of three or more groups, either the Kruskal-Wallis with Dunn’s multiple comparison test or a one-way ANOVA with Tukey-Kramer tests was implemented depending upon the normality of the specific dataset. Statistical analysis performed using Prism 9 and 10 (Graphpad, INC) graphing and statistical software package.

## Notes

### Competing Interest Statement

The authors have declared no competing interest.

